## Supplementary material for "Identification of differentially recognized T cell epitopes in the spectrum of *Mtb* infection": Figure S1 and Table S2

SUPPLEMENTARY Figures and Tables

**Figure S1**

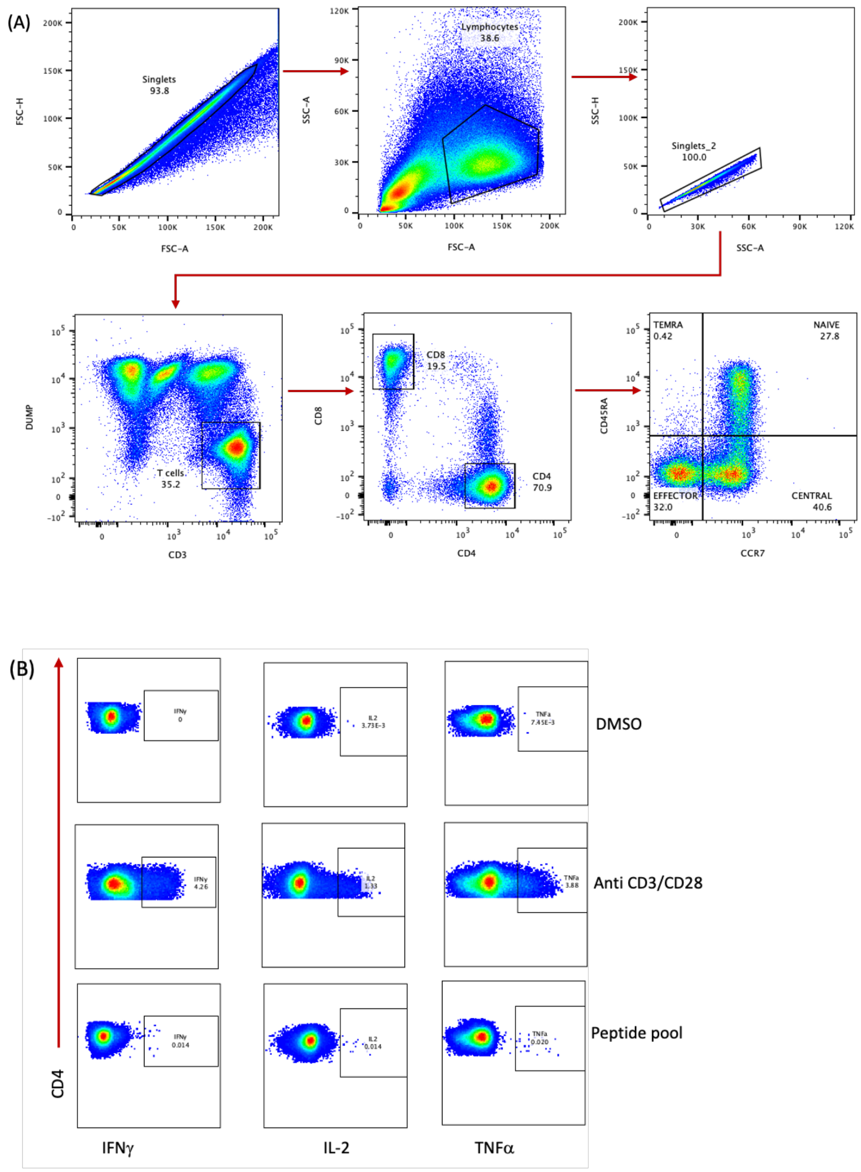


**Figure S1**: Gating strategy for flow cytometry experiments. (A) A representative strategy used to define cytokine-producing T cells. (A)PBMCs were stained and gated on singlets (FSC-A/FSC-H), lymphocytes, singlets (SSC-A/SSC-H), live CD19- CD14- CD3+ T cells, and CD4+ or CD8+ T cells. These were gated on memory populations based on CD45RA and CCR7 expression. (B) Definition of cytokine-producing CD4 T cells stimulated with DMSO (negative control: top), anti-CD3/CD28 (positive control: middle), and peptide pool (MTB300: bottom).

**Table S1. T cell epitopes recognized by 21 participants with ATB (mid-treatment)**
Provided as .xlsx file

**Table S2. Antibodies used for flow cytometry experiments**

| Marker | Clone | RRID | Fluorochrome | Manufacturer | Volume per one million cells |
|---|---|---|---|---|---|
| CD3 | UCHT1 | AB_389310 | AF488 | Biolegend | 2ul |
| CD4 | RPA-T4 | AB_395752 | PE | BD biosciences | 1ul |
| CD8 | RPA-T8 | AB_2874820 | BUV661 | BD biosciences | 1ul |
| CD45RA | HI-100 | AB_10965547 | BV421 | Biolegend | 2ul |
| CD19 | HIB19 | AB_2561668 | BV510 | Biolegend | 1ul |
| CD16 | 3G8 | AB_2562085 | BV510 | Biolegend | 1ul |
| CD20 | 2H7 | AB_2561941 | BV510 | Biolegend | 1ul |
| CD14 | 63D3 | AB_2716229 | BV510 | Biolegend | 1ul |
| CCR4 | 1G1 | AB_396907 | PE-Cy7 | BD biosciences | 1.5ul |
| CCR7 | G043H7 | AB_2563865 | BV711 | Biolegend | 1.5ul |
| CCR6 | 11A9 | AB_2833076 | BUV496 | BD biosciences | 1ul |
| CXCR3 | G025H7 | AB_2563157 | BV605 | Biolegend | 1ul |
| TNF-alpha | MAb11 | AB_2043889 | eF450 | Invitrogen | 1.5ul |
| IFN-gamma | 4S.B3 | AB_469506 | APC | Invitrogen | 1.5ul |
| IL-2 | MQ1-17H12 | AB_2744488 | BB700 | BD biosciences | 2.5ul |
| Live/Dead | 65086614 | | eF506 | Invitrogen | 0.2ul |
